# Allosteric inactivation of an engineered optogenetic GTPase

**DOI:** 10.1101/2022.05.16.490643

**Authors:** Abha Jain, Nikolay V. Dokholyan, Andrew L. Lee

**Author notes:** Andrew L. Lee. **Email:**. **Author Contributions:** A. J., N.V.D., and A.L.L. designed research, A. J. performed research, A.J., and A.L.L. analyzed data, A.J., and A.L.L. wrote the paper.

## Abstract

Optogenetics is a technique for establishing direct spatiotemporal control over molecular function within living cells using light. Light application induces conformational changes within targeted proteins that produce changes in function. One of the applications of optogenetic tools is an allosteric control of proteins via light-sensitive LOV2 domain, which allows direct and robust control of protein function. Computational studies supported by cellular imaging demonstrated that application of light allosterically controlled signaling proteins Vav2, ITSN, and Rac1, but the structural and dynamic basis of such control has yet to be elucidated by experiment. Here, using NMR spectroscopy, we discover principles of action of allosteric control of cell division control protein 42 (CDC42), a small GTPase involved in cell signaling. Both LOV2 and Cdc42 employ flexibility in their function to switch between “dark”/ “lit” or active/inactive states, respectively. By conjoining Cdc42 and LOV2 domains into the bi-switchable fusion Cdc42Lov, application of light – or alternatively, mutation in LOV2 to mimic light absorption – allosterically inhibits Cdc42 downstream signaling. The flow and patterning of allosteric transduction in this flexible system is well-suited to observation by NMR. Close monitoring of the structural and dynamic properties of dark versus lit states of Cdc42Lov revealed lit-induced allosteric perturbations. Chemical shift perturbations for lit mimic, I539E, have distinct regions of sensitivity and both the domains are coupled together leading to bi-directional interdomain signaling. Insights gained from this optoallosteric design will increase our ability to control response sensitivity in future designs.

**Significance Statement:** Control of cell signaling activity in proteins by light is one of the primary goals of optogenetics. The hybrid light-receptor/cell-signaling protein Cdc42Lov was engineered recently as an optogenetic tool, employing a novel allosteric strategy that results in photoinhibition. In contrast to previous activation designs, the mechanism of inhibition of GTPase signaling activity in Cdc42 is only apparent at a detailed structural and dynamic level. NMR characterization of dark and mutationally “lit” forms reveals the allosteric interdomain perturbations, knowledge of which will enhance future applications of this design strategy.

## Introduction

Over the last several years, there has been great interest in controlling the activity of proteins with light (1–8). The field of ‘optogenetics’ has expanded our understanding, *in situ*, of the role of specific proteins in signal transduction, and especially, how neuronal processes are controlled (9–11). In the cellular environment, proteins are naturally controlled in many ways, including allosteric regulation (5, 9–13). Recently, an optogenetic approach was employed to regulate the activity of signal transduction proteins allosterically, by engineering in a modulatory light-sensing domain (6, 14, 15). A computational strategy was employed for introducing the optogenetic sensor– a LOV domain – into loops of proteins that regulate cell motility, which included the two GTPases Cdc42 and Rac1 (6, 16). In the designs, light-induced changes of the LOV domain is transferred into perturbations to the GTPase, resulting in allosteric regulation of activity (6).

Rho family GTPases constitute a family of small (~21 kDa) signaling G proteins that bind and hydrolyze GTP (17). Members of this family, such as Ras, Cdc42, and Rac1, act as molecular switches by shuttling between inactive and active states upon binding to GDP and GTP (17). In the abovementioned study, Cdc42 was engineered by introduction of the *Avena sativa* LOV2 domain such that it is regulatable by light in vivo (6), enabling spatio-temporal control. The general model of LOV2 action is that, upon application of blue light and absorption by FMN cofactor, the LOV2 domain Jα-helix becomes disordered (18, 19). Because the LOV2 domain is at the domain junction and allosterically linked to the active site, the Jα deformation transfers into the Cdc42 domain to down-regulate GTPase activity, resulting in photoinhibition.

There have been several optogenetic designs that couple the LOV2 light switch to signaling proteins (20, 21). Initially, the LOV2 and target proteins were engineered to result in photoactivation, which work by either light-induced dissociation or dimerization of LOV2. While in these cases the activation mechanisms are clear, the allosteric mechanism of photoinhibition (6, 22) is not. Presumably, target domain (e.g. Cdc42) inhibition results from an allosteric transfer of some kind of strain and/or conformational transition, although it is possible that Jα-helix unwinding might lead to target domain unfolding. Obtaining more mechanistic detail into how photoinhibition is achieved will be important to improve the design. A similar question is whether close tethering of the Jα-helix to the target protein in these designs degrade LOV2’s innate allosteric light responsiveness. Ultimately, the major question is how light absorption in LOV2 allosterically propagates into Cdc42, and what structural regions are affected.

To address these questions and gain a first detailed look into an ‘optoallosteric’ protein, we have undertaken a study of the hybrid photoinhibitable protein Cdc42Lov, which to date has only been characterized *in vivo* (6). Cdc42 function was assayed *in vitro* with purified components to obtain a quantitative assessment of optogenetic function, and a detailed NMR analysis has been carried out on Cdc42Lov to assess allosteric mechanism. NMR is a suitable method to probe Cdc42Lov not only because both LOV2 and Cdc42 domains are intrinsically flexible as they are both “switch” proteins, but also because there is likely some interdomain flexibility that may be modulated in the lit state. Both dark and “lit” forms were studied via NMR chemical shift changes, with the “lit” form stabilized through use of a previously characterized LOV2 domain mutation, I539E (23). Extensive NMR chemical shift assignment and TROSY-HSQC spectral changes revealed that overall the Cdc42 domain remains structurally intact in the lit form, and that key functional regions of both domains are perturbed structurally and dynamically, providing insight into the allosteric mechanism. Interestingly, allosteric communication was found to be bidirectional, as binding of Pak1 effector to Cdc42 resulted in significant changes in critical photosensitive regions of LOV2.

## Results and Discussion

The computational design and *in vivo* function of Cdc42Lov, a light-responsive, allosteric fusion of human Cdc42 and the second LOV domain from *Avena sativa* (asLOV2) was described previously (6). In this design, the LOV2 domain is inserted into the β3-β4 loop of Cdc42, such that it is bordered by the N-terminal segment of Cdc42 (1-47) in the preceding residues and the C-terminal segment of Cdc42 (48-178) in the following residues (Figure 1). This mode of insertion positions the LOV2 domain at a different surface from the nucleotide and Cdc42 effector binding surfaces, and results in photoinhibition of Cdc42 activity. *In vivo*, the inhibitory effect was a 2-3 fold activity change for Cdc42Lov, but no characterization has been carried out for single steps *in vitro*, such as how light affects Cdc42 effector binding. Thus, basic mechanistic information on how the Cdc42Lov domain junction partitions and directs destabilizing energy, as well as how this impacts functional interactions, will deepen our understanding of this novel allosteric design, and increase our ability to control response sensitivity in future implementations or new designs. We sought to track the communication pathway linking LOV2 and Cdc42 domains using mutations that mimic light absorption and monitoring with NMR spectroscopy.

**Figure1:**
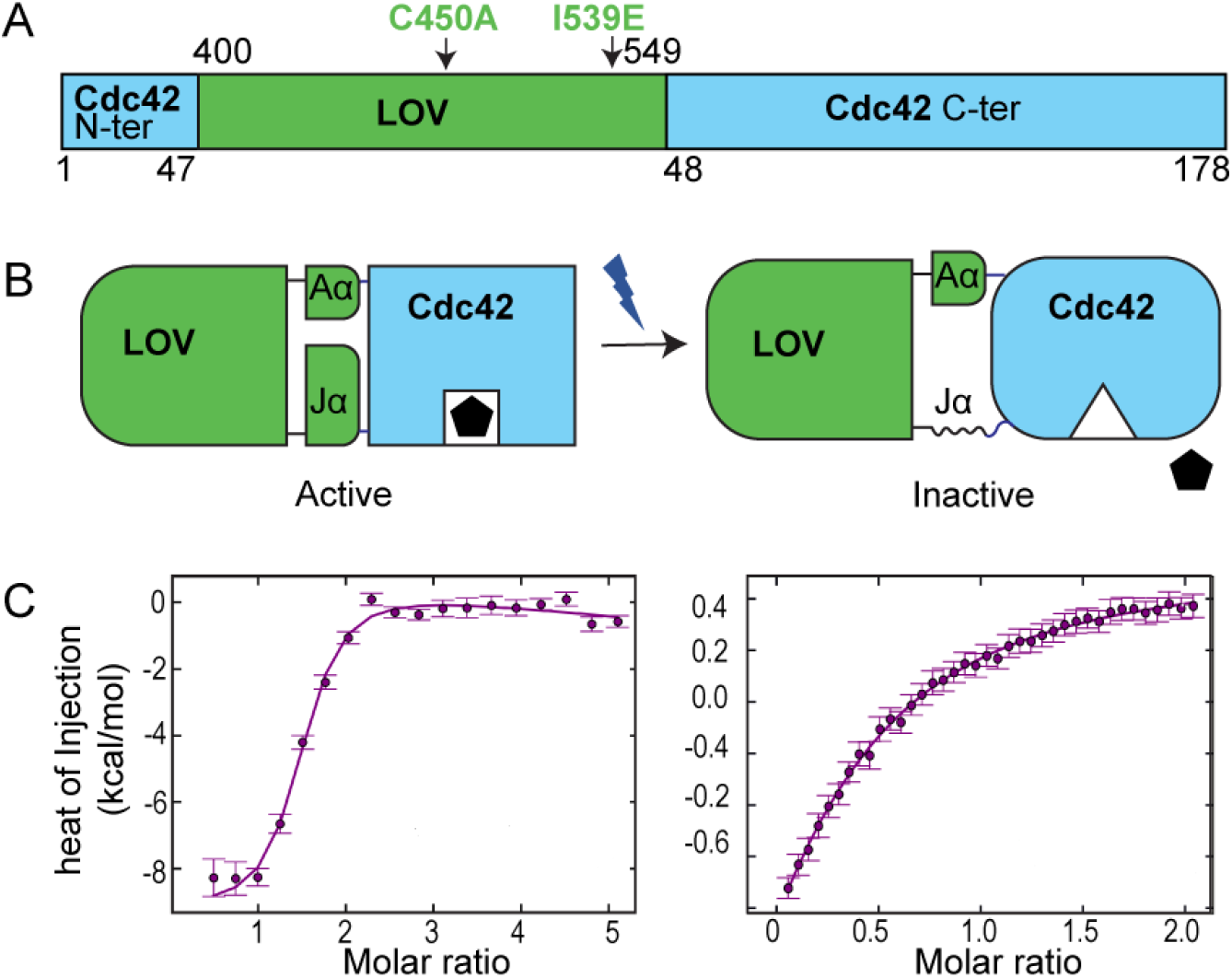
Optogenetic design of allosteric inactivation by light. (A) Gene construct of Cdc42Lov showing insertion of LOV2 domain into Cdc42, ~330 residues. Natural numbering of each domain is retained. On top is LOV2 domain and bottom is Cdc42 numbering. (B) Schematic of allosteric mechanism showing active state changes to an inactive state upon light illumination. Pentagon represents the effector protein/molecule. (C) ITC data for PAK1 effector peptide binding to Cdc42lov^WT, CA^ (left, K_d_ = 0.3 μM) and light-activation mimicking I539E Cdc42lov^LM, CA^ (right, K_d_ = 88 μM).

Several mutations were employed to control states of both domains. For simplicity and relation to familiar residue numbering of Cdc42 and asLOV2, we retain the respective single domain numbering. The I539E (LOV2 domain) mutation mimics the effects of blue light absorption and was, therefore, used as a proxy for the “lit” state of Cdc42Lov (LM) (21). While “dark” state mutants also exist (C450A) (DM), these were typically not needed since measurements on WT protein were made in the absence of light. The Q61L mutation in Cdc42 is known to be constitutively active (24) and was used to favor the GTPase active conformation. GTPase activity is modulated by the exchange of GDP/GTP that toggles inactive vs active conformations (25). In Cdc42Lov, light absorption may alter GDP/GTP exchange or directly alter the effector binding site, or some combination both. Because we found that Cdc42Lov allosteric function exists *in vitro* by using GDP in conjunction with the Q61L mutation (below), NMR studies were conducted with GDP nucleotide, along with the Q61L mutation to favor the active conformation (denoted by CA=constitutively active) as noted (Materials and Methods).

### “Lit” (LM) conformation allosterically inhibits Cdc42 effector binding in vitro

The effect of the I539E lit mutation on Cdc42Lov (Cdc42Lov^LM^) was tested by measuring binding affinity to Cdc42 downstream effector Pak1. Pak1 is a serine/threonine kinase that only binds to the active form of Cdc42 (26), and the interaction can be shown with binding of a ~50-residue Pak1 segment to the constitutively active (CA) Q61L mutant of Cdc42 (27, 28). Pak1 peptide binding was evaluated for Cdc42Lov^WT,CA^ and Cdc42Lov^LM,CA^ using isothermal titration calorimetry (ITC). As expected, the unlit (i.e., WT LOV2 sequence) protein showed high-affinity binding with K_d_ = 0.3 μM, indicating that the Cdc42 domain retains full effector binding function (Figure 1C). By contrast, mutationally “lit” Cdc42Lov^LM,CA^ showed greatly reduced binding affinity, with K_d_ ~ 90 μM (Figure 1C). Thus, *in vitro*, the light-mimicking LOV2 domain perturbation is allosterically transduced to the Cdc42 domain to drastically reduce (> 100-fold) effector binding function.

### NMR spectral comparison of Cdc42Lov^WT^ and Cdc42Lov^LM^

To gain structural insight into the optogenetic allosteric mechanism of Cdc42Lov, NMR signals of “lit” Cdc42Lov^LM^ were compared to “unlit” Cdc42Lov^WT^. We confirmed that the 2D ^1^H-^15^N HSQC spectrum of the wild-type protein was nearly identical compared to Cdc42Lov^DM^ (Figure S1) due to the absence of visible light in the magnet. While both lit and unlit forms show spectra with numerous peaks and significant chemical shift dispersion indicative of folded proteins, the unlit (WT) form has more peaks with uniform intensity (Figure 2). Spectral overlay suggest the presence of some slow μs-ms dynamics in the lit state. The greater degree of variability in peak intensities, along with some peaks collapsed into the center region, in the lit form spectrum suggests a subpopulation of destabilized species or regions of partially unfolded structure in the lit form (Figure 2). Interestingly, circular dichroism (CD) spectra of both forms showed only a slight difference (Figure S2), indicating that the lit form appears to retain nearly all of its helical structure, and SEC-MALS also showed similar sizes for lit and unlit states (Figure S2). Given that the gross structural features of lit and unlit Cdc42Lov appear similar, the NMR signals were analyzed in greater detail to detect differences in the two forms.

**Figure 2:**
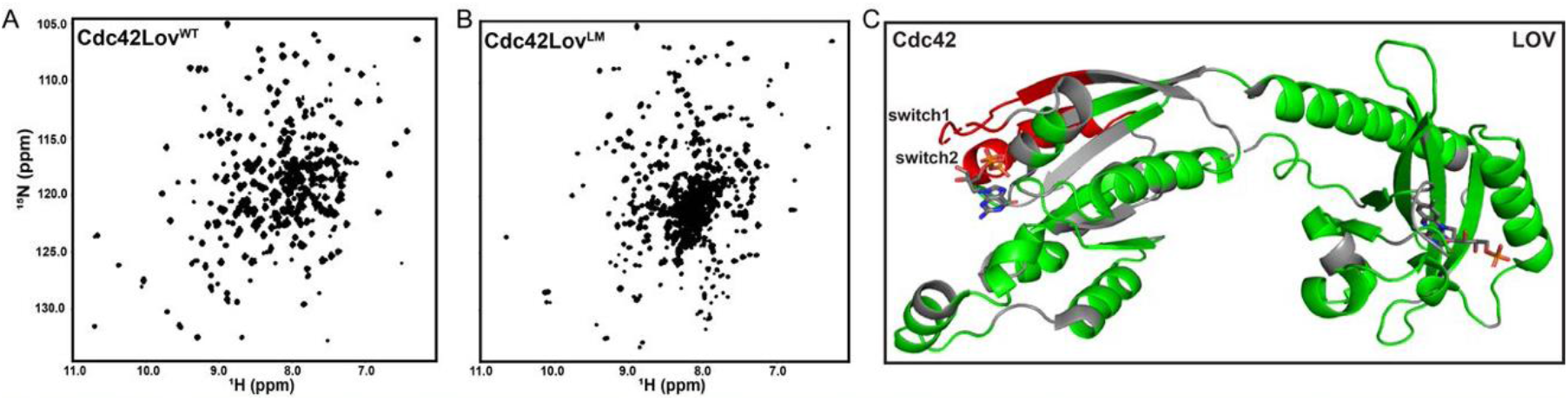
TROSY-HSQC of unlit and lit state of Cdc42Lov design. (A) Cdc42Lov^WT^ “unlit” HSQC (B) Cdc42Lov^LM^ “lit” (C) Backbone assignment of Cdc42Lov^WT^. Assignments are shown on a model generated using the SWISS modeler. In green are assigned residues, grey unassigned, and in red are the unassigned switch regions of the Cdc42 domain.

### Light-triggered interdomain force transduction monitored by chemical shift perturbations

To gain structural insight into how light allosterically transmits from LOV2 to Cdc42, chemical shift perturbations (CSPs) were recorded. As the design couples initial LOV2 domain perturbation to the Cdc42 domain, significant CSPs in the LOV2 domain, especially from Jα-helix unwinding, are expected. Backbone chemical shifts for Cdc42Lov^WT^ and Cdc42Lov^LM^ were assigned using standard triple-resonance methods on perdeuterated proteins (Methods and Materials). Assignments for unlit and lit forms were made for 68% and 62% of non-proline residues, respectively, with peak broadening from dynamic behavior being the primary reason for unassigned residues (Figure 2C). In principle, CSPs can give insights into adjustments made in both domains and trace critical residues that lie along the communication pathway between LOV2 and Cdc42 functional sites (Figure 3B, S3). Lit form-induced chemical shift effects are shown in Figure 3. One of the key LOV2 domain residues is C450, which is known to form a covalent adduct with FMN upon illumination (28). The unlit state has a prominent amide peak for C450 which becomes completely broadened in Cdc42Lov^LM^ (Figure 4a). Similarly, Q513, which serves as a “glutamine lever” that induces unwinding of the Jα-helix (Figure 4b), is also completely broadened in Cdc42Lov^LM^ (29). Adduct formation and glutamine lever movement are coupled events, and their broadening suggests that the coupling is retained in Cdc42Lov^LM^. Overall, this behavior in the LOV2 domain is fully consistent with I539E serving as an effective proxy for light absorption.

**Figure 3:**
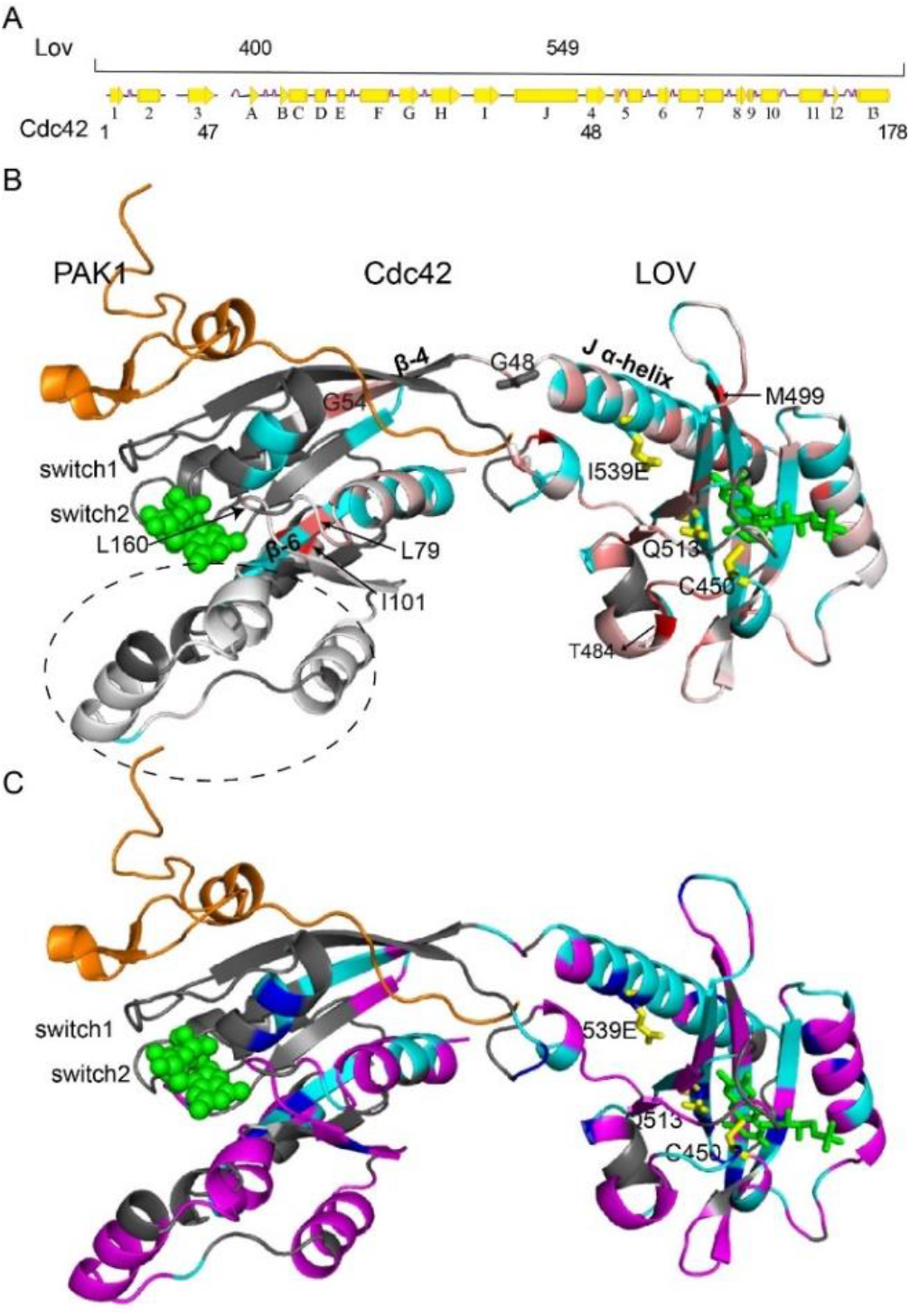
Chemical shift perturbation map. (A) Secondary structure for Cdc42Lov and the numbering for each domain. (B) Perturbation map shown in white-to-red scale showing increasingly perturbed residues. Broadened residues are in cyan, and unassigned residues are in black. (C) Intact-residue map based on the intensity of peaks. Intact residues are highlighted in magenta, cyan are residues with non-zero CSPs, blue are completely broadened residues, and unassigned residues are in black. FMN and GDP are shown in green spheres and Pak1 (PDB1e0a) is shown in orange. Side chain of residues involved in signal transduction for LOV2 are shown in yellow sticks.

**Figure 4:**
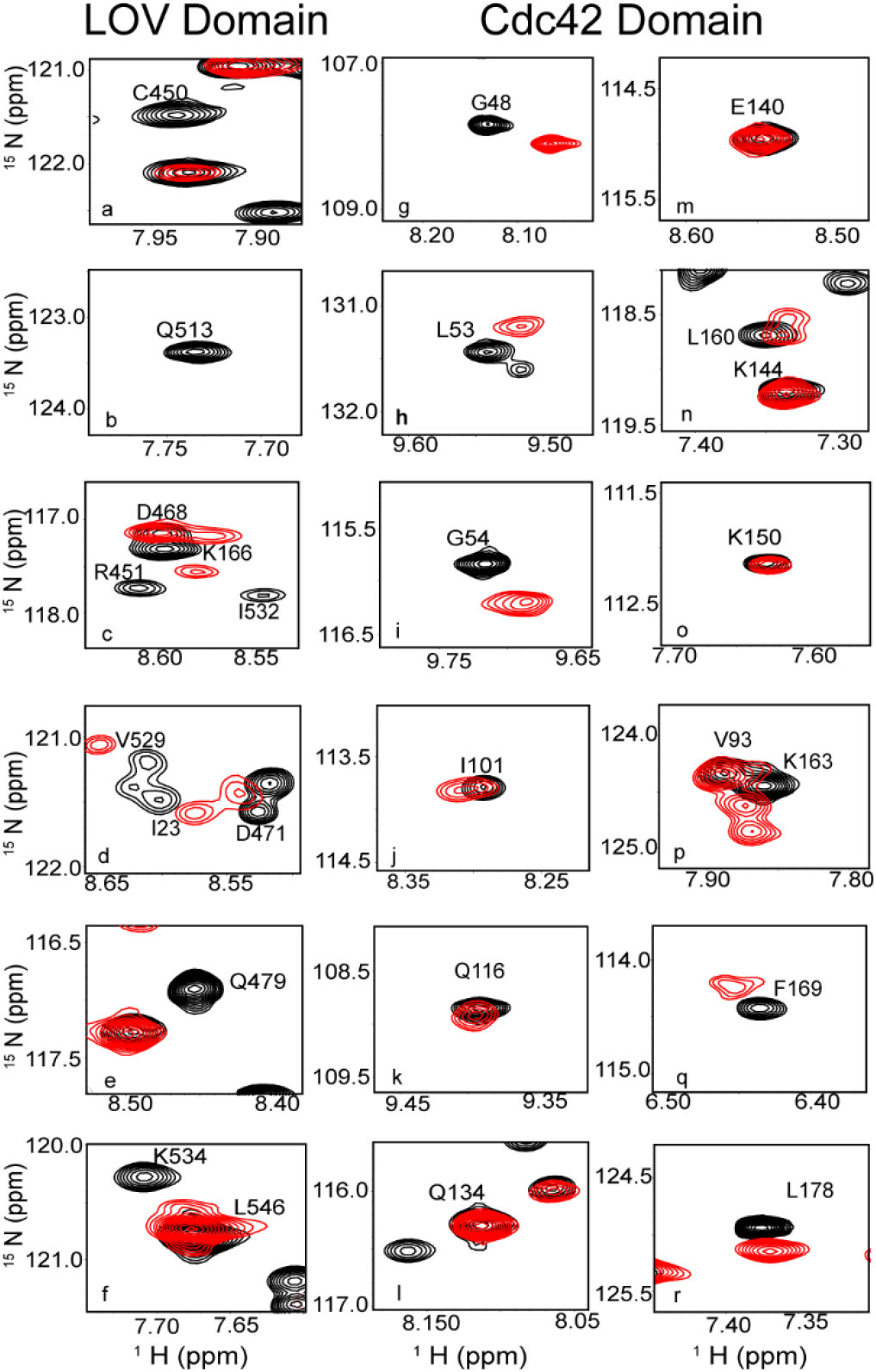
CSP peak comparison of unlit and lit states. Cdc42Lov^WT^ “unlit” peaks are shown in black and Cdc42Lov^LM^ I539E “lit” mimic peaks are in red.

In line with the coupling design, the I539E mutation results in extensive CSPs and peak broadening for various secondary structures surrounding the FMN binding pocket. This includes residues in β-strands G, H, and I, such as R451, I466, D471, V478, Q479, F494, L496, M499, Y508, F509, and L514 (Figure4, S4) with these residues showing either peak shifting or disappearance/broadening, indicative of a structural rearrangement. Although light activation leads to the unwinding of the Jα-helix in native LOV2 ^6,28^, strict CSPs do not indicate complete unwinding of the Jα-helix in Cdc42Lov ^LM^. This is likely because CSPs are based entirely on the peak position of assigned residues, and only require that peaks be present and assignable. Closer inspection showed that some assigned peaks in the LOV2 domain are greatly reduced in their intensity (or broadened). Thus, while CSPs can track perturbations, they can be less sensitive to changes in dynamics or local stability.

To address this caveat, we included another metric for perturbation based on intensities. Here the residues are binned as “intact” if the peak position and intensity are comparable to the unlit form, or else it is identified as “not intact”. This intact-residue map (Figure 3C) provides another level of understanding for lit-form perturbations and shows which protein regions retain stably folded structure. The core region with FMN binding pocket shows an equal mixture of intact and non-intact residues (Figures 3C, S5). Almost all the residues of the Jα-helix including I539 appear as non-intact except L546 (Figure 4, S5) and G547, part of the hinge region, and L531. Together, CSPs and the intactness map show that although the Jα-helix in the LOV2 domain appears to have some residues that are minimally perturbed, the entire Jα-helix is undergoing transitions or conformational change.

Interestingly, HSQC peak perturbations are also observed in the Cdc42 domain of Cdc42Lov^LM^. These perturbations are somewhat weaker compared to those in the LOV2 domain but nevertheless indicate that Cdc42Lov^LM^ is transducing the light-induced effect into Cdc42. A major region of interest is the Pak1 binding interface since application of light alters Pak1 binding (Figure 1C) (27). Residues in Cdc42 that interact with Pak1 are 21-25 (α2), 36-47 (β3), 67-72 (switch 2 or α5), and 166-178 (α13). Of these residues, approximately half were assigned and were used to monitor the impact of I539E/”light” perturbation. The majority of these were observed in the C-terminal segment, within α13, by either exhibiting CSPs or broadening (Figure 3, S6). Specifically, residues K166, D170, I173, and L174 are impacted by the I539E mutation and thus are likely involved in allosterically reducing Pak1 binding affinity (Figure 4c). While most of the N-terminal segment of the Cdc42 domain could not be assigned, there was notable broadening in a few residues within α2, including I23 (Figure 4d) which makes extensive contacts with F81 of Pak1. It is likely that switches 1 and 2 are also perturbed by I539E, but these regions could not be assigned in any state, which is typical for Cdc42 and GTPases in general (30)

To get better insight into how light-induced signaling propagates into Cdc42 and effector binding regions, we looked at the connecting domain hinge region and other distal parts of Cdc42. At the hinge, between loops β3-β4, a few residues, L546, G547 (both in LOV2), and G48 (Cdc42), are slightly perturbed but still appear intact, suggestive of the strong connection between the two domains. Furthermore, the CSPs in the Jα-helix propagate to the adjacent β4 strand of the Cdc42 domain (Figure 3B). T52, L53, and G54 are a few of the β4 strand residues that experience perturbation due to the unwinding of Jα-helix. Thus, Jα-helix unwinding is translating structural changes to the connected β4 strand. However, light-induced changes extend beyond the immediately connected strands to the Jα-helix. For example, residues in the β6 strand (F78-F82) show peak shifting or broadening (Figure 3B). F78 and V80 makes a triad with distal residue I101, that also experiences a high degree of light-induced chemical shift perturbation (Figure 3B, 4). Except for L79, all other residues in the β6 strand are broadened, confirming that the I539E mimic of light absorption penetrates central regions of Cdc42.

The other primary region for observing structural perturbations is the nucleotide-binding site because of the functional coupling between nucleotide status (GDP vs. GTP) and effector binding in GTPases. In Cdc42, the nucleotide-binding site is formed by the switch regions switch 1 (28-40) and switch 2 (57-74) (27, 28), and the three loops formed by residues 12-17 (connecting β1-α2), 115-120, and 158-161 (connecting β12-α13) (Figure S6). Perturbations to the switch regions may be expected via translation through the connecting strands β3 and β4. However, because of the intrinsic dynamic nature (loss of NMR signals) of the two switches, we could only observe perturbations to the three loops. Specifically, L160 shows a sizeable shift perturbation (0.05 ppm) (Figure S6), and T115 is broadened, indicating a mixture of structural perturbation and conformational instability in the nucleotide-binding site upon I539E mutation (Figure 3B).

To our surprise, the core region of the Cdc42 domain, Q116-K150, shows little perturbation and remains intact, suggestive of controlled transmission with directed allosteric communication (i.e., some regions are entirely unperturbed) from the LOV2 domain (Figure 3B, 3C, 4). This unperturbed intact core region is followed by a C-terminal helix (α13) that showed a high magnitude of light dependent chemical shift perturbation suggesting an important role in signal propagation from either nucleotide-binding site to effector binding site or vice-versa.

Collectively, these HSQC peak perturbations show that Cdc42Lov ^LM^ is undergoing allosteric conformational change, with both domains showing perturbation in the lit-mimicked state. Both peak shifting (CSPs) and broadening (intactness) suggest that the LOV2 domain is highly perturbed, and residues around the FMN binding pocket are undergoing conformational change, and the Jα-helix largely unwinds in the lit state. This Jα-helix unwinding is further transmitted to the Cdc42 domain, mainly around β4 and β6, leading to CSPs. It suffices to say that conformational change in the Cdc42 domain is mainly around regions that are in close proximity to the nucleotide-binding region and C-terminal helix. Also, there is a structurally unperturbed intact region suggestive of controlled transmission of signals. Overall, the data support that this Cdc42Lov ^LM^ mimic, an observed “lit” state, is working according to the strategy of inserting the LOV2 domain.

### Pak1 bound CSP’s confirming bi-directional allosteric communication

To further evaluate the coupling of domains, we observed how Pak1 binding (downstream effector) affects Cdc42Lov. Since Pak1 only binds to the active state of Cdc42, we used Cdc42Lov ^WT,CA^ (Q61L, constitutively active) to study the complex. Upon binding Pak1, substantial chemical shift perturbations were observed in α2 (K18-T27), α7 (F90-E95), α10 (D122-K135), and α11(A142-D148) in the Cdc42 domain (Figure 5, S7), all of which have been shown to undergo conformational changes upon binding Pak1 (24, 31, 32) (Figure 5, S7). In addition to the above helices, extensive broadening and perturbations were observed in the α13 (L165-E178) helix (Figure S8). Interestingly, α13 is also perturbed in I539E lit mimic, suggesting a possible coupled behavior between both domains.

**Figure 5:**
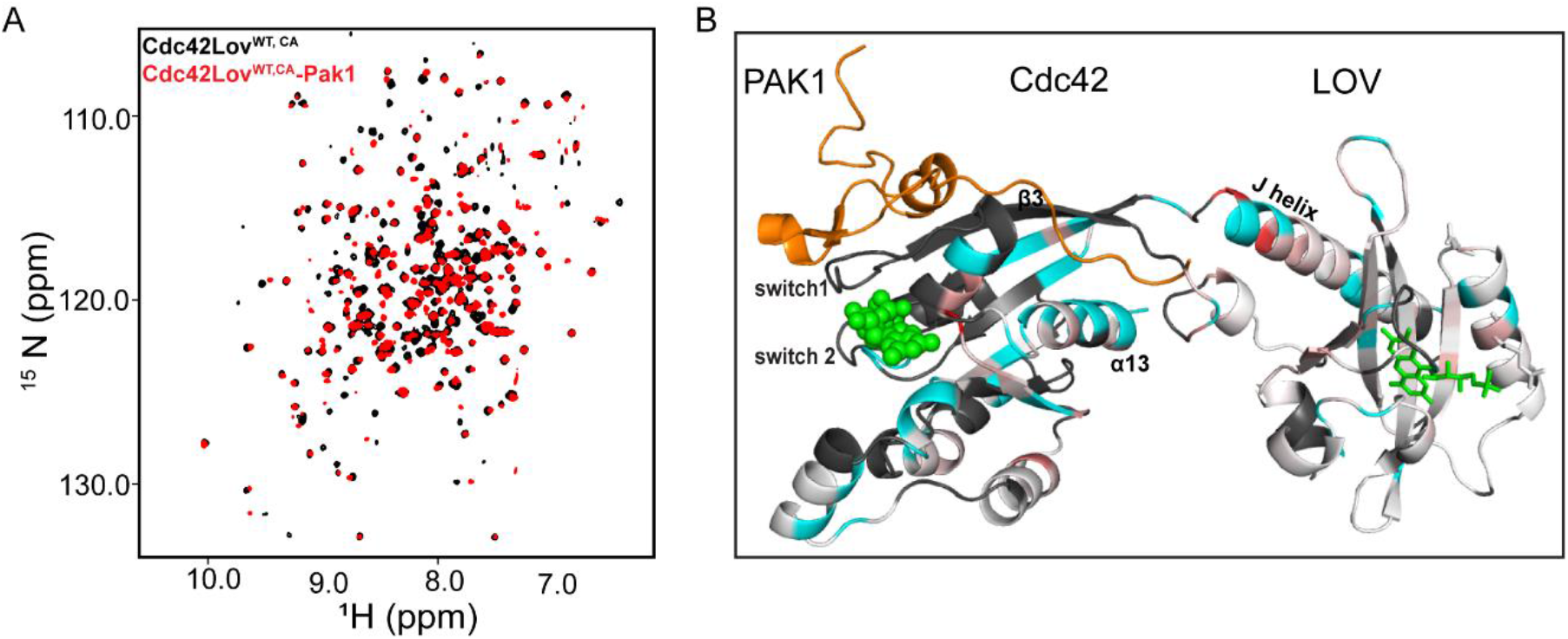
Pak binding (A) Overlay of ^15^N TROSY-HSQC of ^CA^Cdc42Lov^WT^ and ^CA^Cdc42Lov^WT^ Pak1 bound in black and red respectively. (B) Chemical shift perturbation map for Pak1 binding to Cdc42Lov^WT^. Color coding as in Figure 3B.

To our surprise, Pak1 binding introduced changes not only in the Cdc42 domain but also in the LOV2 domain (Figure 5, S7). Overall perturbations in the LOV2 domain are in the core region and around the Jα-helix. Closer inspection showed that R451, residue next to C450, shows chemical shift changes upon Pak1 binding, indicative of allosteric communication to the key residue involved in adduct formation in the lit state. Similarly, residues close to the glutamine (Q513) lever, I510 and G511, were found perturbed upon Pak1 binding. These were the sites responsive to I539E (light-mimic), yet it is striking that these same sites are clearly perturbed upon distal binding of Pak1. Similarly, in the Cdc42 domain, I101, L160, and a majority of the α13 helix were all perturbed by both I539E mutation and Pak1 binding. Taken together, these in-common sites of perturbation indicate that the nature of domain coupling leads to bidirectional interdomain signaling (Figure S9).

In this study, we have tracked the allosteric response of an optogenetically encoded structural distortion in an engineered two-domain hybrid protein, Cdc42Lov. Though the allosteric photoinhibition of Cdc42 function was consistent with the design, knowledge of the operative mechanism has been lacking. Due to the flexible nature of the interdomain linkage, we have opted for a solution-based approach to reveal clues into the allosteric mechanism. NMR characterization of the light-mimicking response of I539E has the advantage of site-specific tracking using chemical shifts and peak intensities that report on dynamics. Monitoring of chemical shifts allowed regions in Cdc42Lov to be identified as experiencing structural perturbation and dynamic switching behavior – as well as general regions of the protein that remain structurally intact vs. locally destabilized – in response to light absorption. As protein engineering advances to adopt similar, improved, or alternative strategies to implement allosteric functional properties, the high-resolution solution approach afforded by NMR should be a valuable tool for assessing such mechanisms, especially when systems possess flexibility that may make other structural approaches more challenging.

## Materials and Methods

### Protein Expression and Purification

The photosensitive construct of Cdc42 using LOV2 insertion (Cdc42Lov, Fig. 1) was designed and synthesized as described previously (6) (Bio basic inc.) with an often-used Q61L mutation that confers constitutively active Cdc42 (Cdc42CA). The C450A and I539E, “dark” and “lit” mutants, as well as Pak1 binding protein (Pak1) domain constructs with a C-terminus 6xHis-tag (PET23-PBP(65-109)-N-Cys-His6) were expressed in *E. coli* and purified as described in SI Methods.

### Isothermal Titration Calorimetry

ITC samples for all variants and Pak1 peptide were prepared in NMR buffer with TCEP as a reducing agent at pH 7.5. Experiments were conducted on a MicroCal AutoITC 200 (Malvern Panalytical, Malvern, UK). Protein concentration in the cell was set to 20 and 150 μM, for unlit and lit respectively. Pak1 concentration was10 times higher depending upon cell protein concentration. Raw thermograms were integrated using NITPIC (33) and fitted in SEDPHAT (34) software respectively. The single-site binding model was used for data fitting.

### NMR spectroscopy

Standard transverse relaxation-optimized spectroscopy (TROSY) triple resonance and ^1^H-^15^N heteronuclear single quantum coherence (TROSY-HSQC) experiments were used for backbone assignment and CSP calculations(35), respectively, as described in SI Methods.

## Acknowledgments

The work was supported by National Institute of Health Grants 1R35 GM134864 and 1RF1AG071675 (to N.V.D.) and GM083059 (to A.L.L.). N.V.D. also acknowledges the support from the Passan Foundation. The UNC School of Medicine Biomolecular NMR Lab is supported by the National Cancer Institute of the National Institutes of Health under award number P30CA016086. We thank Dr. Ashutosh Tripathy of the UNC Macromolecular Interactions Facility for assistance in ITC, CD, and SEC-MALS data collection.

## References

1. A. Bansal, S. Shikha, Y. Zhang, Towards translational optogenetics. Nature Biomedical Engineering, 1–21 (2022).

2. L.-D. Huang, Brighten the Future: Photobiomodulation and Optogenetics. Focus 20, 36–44 (2022).

3. P. P. Prosseda, M. Tran, T. Kowal, B. Wang, Y. Sun, Advances in Ophthalmic Optogenetics: Approaches and Applications. Biomolecules 12, 269 (2022).

4. P. Tan, L. He, Y. Huang, Y. Zhou, Optophysiology: Illuminating cell physiology with optogenetics. Physiological Reviews (2022).

5. N. V. Dokholyan, Controlling allosteric networks in proteins. Chemical reviews 116, 6463–6487 (2016).

6. O. Dagliyan et al., Engineering extrinsic disorder to control protein activity in living cells. Science 354, 1441–1444 (2016).

7. N. V. Dokholyan, Nanoscale programming of cellular and physiological phenotypes: inorganic meets organic programming. NPJ systems biology and applications 7, 1–5 (2021).

8. Y. L. Vishweshwaraiah, J. Chen, V. R. Chirasani, E. D. Tabdanov, N. V. Dokholyan, Two-input protein logic gate for computation in living cells. Nature communications 12, 1–12 (2021).

9. C. P. O’Banion, D. S. Lawrence, Optogenetics: a primer for chemists. ChemBioChem 19, 1201–1216 (2018).

10. I.-W. Chen, E. Papagiakoumou, V. Emiliani, Towards circuit optogenetics. Current opinion in neurobiology 50, 179–189 (2018).

11. S. Beck et al., Synthetic light-activated ion channels for optogenetic activation and inhibition. Frontiers in neuroscience 12, 643 (2018).

12. S. J. Wodak et al., Allostery in its many disguises: from theory to applications. Structure 27, 566–578 (2019).

13. J. Wang et al., Mapping allosteric communications within individual proteins. Nature communications 11, 1–13 (2020).

14. O. Dagliyan, K. M. Hahn, Controlling protein conformation with light. Current opinion in structural biology 57, 17–22 (2019).

15. J. Chen, Y. L. Vishweshwaraiah, N. V. Dokholyan, Design and engineering of allosteric communications in proteins. Current Opinion in Structural Biology 73, 102334 (2022).

16. O. Dagliyan, N. V. Dokholyan, K. M. Hahn, Engineering proteins for allosteric control by light or ligands. Nature protocols 14, 1863–1883 (2019).

17. A. Hall, Rho family gtpases. Biochemical Society Transactions 40, 1378–1382 (2012).

18. A. I. Nash, W.-H. Ko, S. M. Harper, K. H. Gardner, A conserved glutamine plays a central role in LOV domain signal transmission and its duration. Biochemistry 47, 13842–13849 (2008).

19. E. Peter, B. Dick, S. A. Baeurle, Mechanism of signal transduction of the LOV2-Jα photosensor from Avena sativa. Nature communications 1, 1–7 (2010).

20. A. Winkler et al., Structural details of light activation of the LOV2-based photoswitch PA-Rac1. ACS chemical biology 10, 502–509 (2015).

21. L. He et al., Circularly permuted LOV2 as a modular photoswitch for optogenetic engineering. Nature Chemical Biology 17, 915–923 (2021).

22. D. Strickland, K. Moffat, T. R. Sosnick, Light-activated DNA binding in a designed allosteric protein. Proceedings of the National Academy of Sciences 105, 10709–10714 (2008).

23. S. M. Harper, J. M. Christie, K. H. Gardner, Disruption of the LOV-Jα helix interaction activates phototropin kinase activity. Biochemistry 43, 16184–16192 (2004).

24. D. I. Johnson, Cdc42: an essential Rho-type GTPase controlling eukaryotic cell polarity. Microbiology and Molecular Biology Reviews 63, 54–105 (1999).

25. R. G. Hodge, A. J. Ridley, Regulating Rho GTPases and their regulators. Nature reviews Molecular cell biology 17, 496–510 (2016).

26. U. G. Knaus, G. M. Bokoch, The p21rac/cdc42-activated kinases (paks). The international journal of biochemistry & cell biology 30, 857–862 (1998).

27. W. K. Stevens et al., Conformation of a Cdc42/Rac interactive binding peptide in complex with Cdc42 and analysis of the binding interface. Biochemistry 38, 5968–5975 (1999).

28. S. M. Harper, L. C. Neil, K. H. Gardner, Structural basis of a phototropin light switch. Science 301, 1541–1544 (2003).

29. J. N. Iuliano et al., Unraveling the mechanism of a LOV domain optogenetic sensor: A glutamine lever induces unfolding of the Jα helix. ACS chemical biology 15, 2752–2765 (2020).

30. H. R. Mott, D. Owen, Allostery and dynamics in small G proteins. Biochemical Society Transactions 46, 1333–1343 (2018).

31. J. L. Feltham et al., Definition of the switch surface in the solution structure of Cdc42Hs. Biochemistry 36, 8755–8766 (1997).

32. D. Gizachew, W. Guo, K. K. Chohan, M. J. Sutcliffe, R. E. Oswald, Structure of the complex of Cdc42Hs with a peptide derived from P-21 activated kinase. Biochemistry 39, 3963–3971 (2000).

33. S. Keller et al., High-precision isothermal titration calorimetry with automated peak-shape analysis. Analytical chemistry 84, 5066–5073 (2012).

34. H. Zhao, G. Piszczek, P. Schuck, SEDPHAT-a platform for global ITC analysis and global multi-method analysis of molecular interactions. Methods 76, 137–148 (2015).

35. A. L. Lee et al., Chemical shift mapping of the RNA-binding interface of the multiple-RBD protein sex-lethal. Biochemistry 36, 14306–14317 (1997).

