## Supplemental text for "Allosteric inactivation of an engineered optogenetic GTPase"

[a]Division of Chemical Biology and Medicinal Chemistry, UNC Eshelman School of Pharmacy, University of North Carolina at Chapel Hill, Chapel Hill, NC 27599
[b] Departments of Pharmacology, and Biochemistry & Molecular Biology, Penn State College of Medicine, PA 17033
[c] Department of Biochemistry and Biophysics, School of medicine, University of North Carolina at Chapel Hill, Chapel Hill, NC, 27599
* Andrew L. Lee.

**The Supplementary information contains SI Materials and Methods and Supplementary Figures S1-S9**

#### Materials and Methods

##### Cdc42Lov construct

The engineered construct of Cdc42 using LOV2 insertion (Cdc42Lov, Fig. 1) was synthesized with Q61L constitutively active mutation (Bio Basic INC). The synthesized gene was inserted between the NdeI and XhoI sites of pET24a, following the N-terminal 6xHis tag, and transformed into BL21 (DE3) star cells. The C450A and I539E, "dark" and "lit" mutants, as well as other non-constitutive mutants, were made using the QuickChange (Agilent) protocol.

##### Protein expression and purification

For the preparation of unlabeled protein for ITC, CD, and SecMALs studies, proteins were expressed in *E. coli* strain BL21 Star (DE3) in 50ml LB media overnight and were used to inoculate one liter of LB. At an $OD_{600}$ of 0.8-1.0, expression was induced with 1 mM isopropyl β-D-1-thiogalactopyranoside (IPTG), and the growth was continued at 18°C for 20 hrs. For isotope-labeled, perdeuterated NMR samples, proteins were expressed in 1 L of M9 (99.8% $D_2O$; Cambridge Isotope Laboratories (CIL)) media with 1 g of $^{15}NH_4Cl$ (CIL) and 2 g of either U-[$^2$H]-glucose or U-[$^{13}$C, $^2$H]-glucose (CIL) as the sole nitrogen and carbon sources, respectively. Cells were harvested by centrifugation and the pellet was resuspended in buffer A (20 mM Tris-HCl, 500 mM NaCl, 25 mM imidazole, 0.5 mM TCEP, 5 mM MgCL2, and 0.10% $NaN_3$ pH 7.5) along with the 200 μM nucleotide (GDP/GMP-PNP), one protease inhibitor tablet, EDTA (ethylenediamine tetraacetic acid) free (Thermo scientific), and lysozyme. Protein purification was performed as described previously (6) with slight modification. Briefly, 6xHis-tagged protein was run over a Ni column (HisTrap HP 5 mL; GE Healthcare, Chicago, IL) using an ÄKTA purifier fast protein liquid chromatography system (GE Healthcare) and tagged Cdc42Lov was eluted using the resuspension buffer A with 500 mM imidazole. The 6xHis-tag was cleaved by incubation with tobacco etch virus (TEV) protease overnight at 4°C. The cleaved protein was subsequently run over the Ni column and collected in the flow through. Untagged protein was concentrated to 5 mL and run over a HiLoad Superdex 200 column (GE Healthcare) in NMR buffer (25 mM Tris-Cl, 150 mM NaCl, 5 mM $MgCl_2$, 0.10% $NaN_3$, and 5 mM DTT pH 7.5). The purified protein was further concentrated to 350 μM and used for ITC or NMR studies.

Pak1 binding protein (Pak1) domain constructs with a C-terminal 6xHis-tag (pET23-PBD(65-109)-N-Cys-His6) was a gift from Klaus Hahn at UNC (Addgene). This construct was transformed into BL21 (DE3) star cells. Protein was expressed as above with the exception that after 1 mM IPTG induction cell growth was allowed for 5 hours at 18°C. Cells were lysed using French press to increase the soluble yield. 6xHis-tag based peptide purification was performed using the same buffer composition described above and passed over a HiLoad Superdex 200 column (GE Healthcare) to obtain high purity.

**NMR spectroscopy**

For NMR experiments on different mutants of Cdc42Lov TROSY-HSQC (transverse relaxation optimized spectroscopy- heteronuclear single quantum coherence), spectra were collected on 850 or 600 MHz Bruker Avance III spectrometers (Bruker, Billerica, MA) equipped with TCI cryoprobes. All datasets were acquired using the trosyf3gpphsi19.2 Bruker pulse program at 298 K.

For backbone resonance assignment of $Cdc42Lov^{WT,CA}$, 350 uM protein sample of U-[$^{2}H$, $^{13}C$, $^{15}N$] was prepared with 5% $D_2O$ and 1 mM GDP in NMR buffer. Six sets of TROSY triple resonance experiments were acquired on 850 or 600 MHz at 298 K: HN(CA)CB, HNCA, HNCO, HN(CA)CO, HN(CO)CA, and HN(COCA)CB.

Because of near-perfect HSQC overlap, resonances for Cdc42Lov$^{WT}$ were assigned based on peak proximity to Cdc42Lov$^{WT,CA}$. Since the sample life of Cdc42Lov$^{LM}$ was less than 10 hours, we used a similar approach for assigning the Cdc42Lov$^{LM}$ where resonances were assigned using peak proximity to Cdc42Lov$^{WT}$. For Cdc42Lov$^{LM}$ we used information from data sets collected at 600 MHz and 800 MHz to confirm peak positions for all assigned residue. This information helped in confirming assignments for more than 100 assigned peaks.

**Calculations of chemical shift perturbations**

Chemical shift perturbations (CSPs, in ppm) for Cdc42Lov$^{LM}$ and Cdc42Lov$^{WT\text{-}Pak1}$ bound states were calculated using a linear relationship based on the distance between amide peak as observed in TROSY-HSQC  for each residue to the nearest peak (minimal shift) in Cdc42Lov$^{WT}$.

$$\mathrm{CSP} = \sqrt{\left(\mathrm{H}(i) - \mathrm{H}'(i)\right)^2 + 0.15 * \left(N(i) - N'(i)\right)^2}$$

where H($i$) is the $_1$H chemical shift of unlit state; H'($i$) is the $_1$H chemical shift of "lit" or peptide; N($i$) is the $_{15}$N chemical shift of unlit state; N'($i$) is the $_{15}$N chemical shift of "lit" or peptide.

### Supplementary Figures

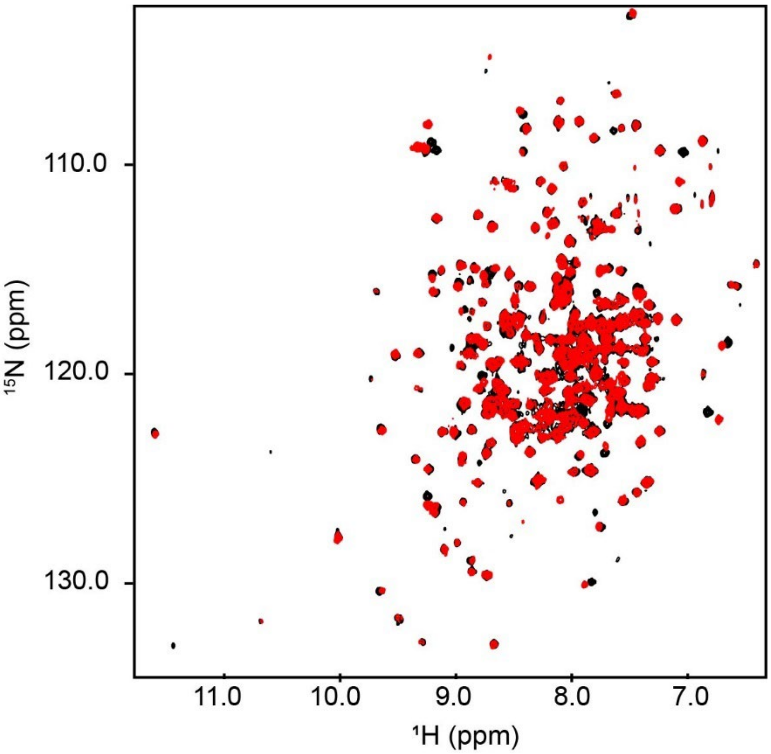


**Figure S1.** Overlay of $^{15}$N TROSY-HSQC of Cdc42Lov$^{WT,CA}$ (black) and Cdc42Lov$^{DM,CA}$ (red).

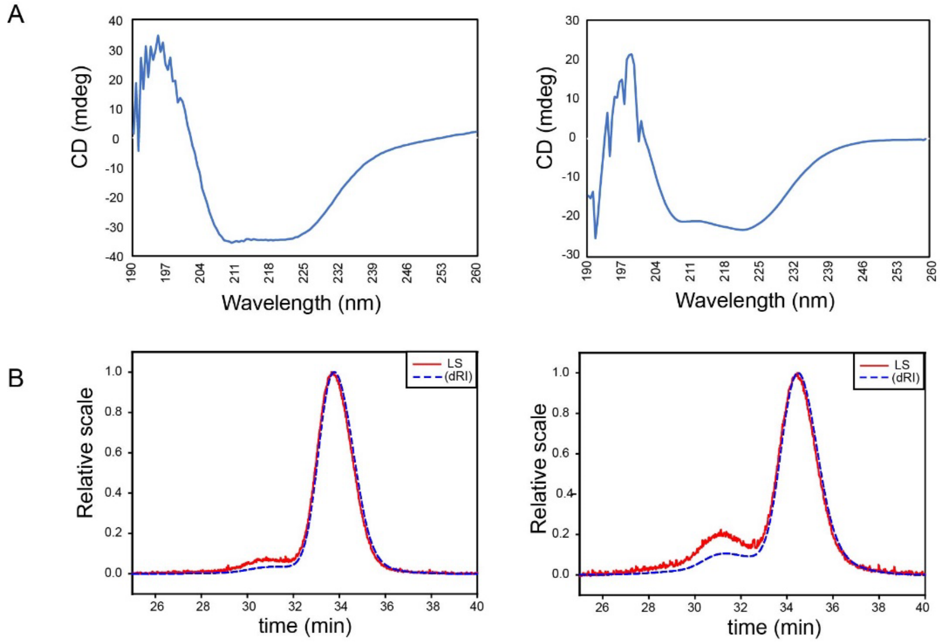


**Figure S2.** Secondary structure and stability of Cdc42Lov$^{WT,CA}$ (unlit) and Cdc42Lov$^{LM,CA}$ (lit). (A) Circular dichroism spectra of unlit (WT, left) and lit (LM, right) state. (B) SEC-MALS peaks indicate similar mass and size of both unlit and lit states, respectively.

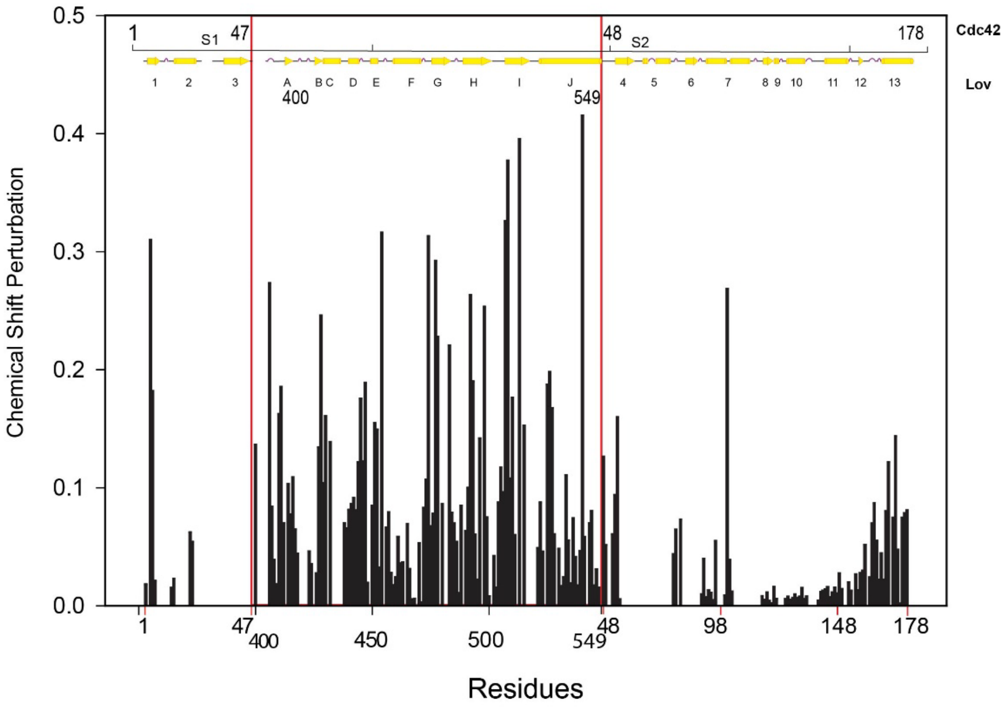


**Figure S3.** CSPs for I539E 'lit' mimic for each assigned peak of Cdc42Lov.

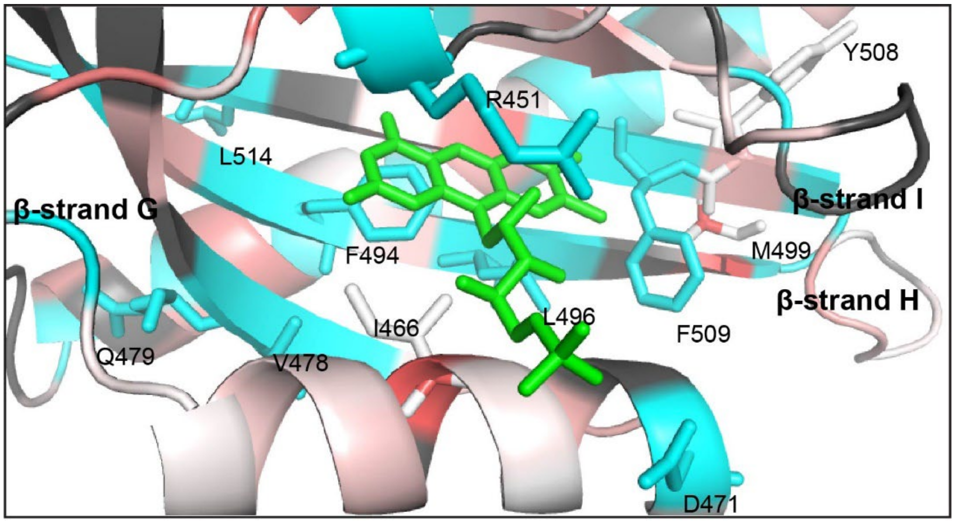


**Figure S4.** CSPs for FMN binding region. The perturbation map is shown in white to red scale. Residues highly perturbed are in red whereas less perturbed residues are shown in white. In cyan are broadened residues. Unassigned residues are in black. FMN is shown as green sticks. A few residues that are perturbed are shown in sticks and labeled.

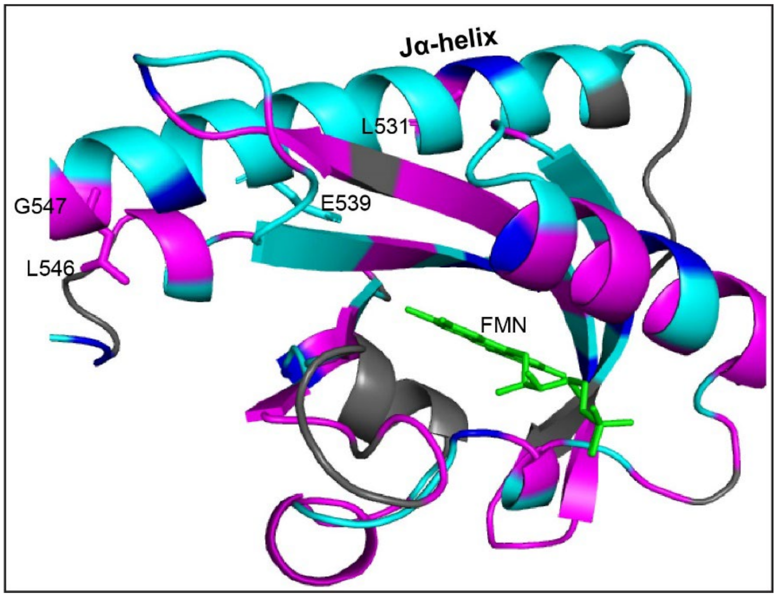


**Figure S5.** Intact residue map based on peak intensity for the FMN binding pocket, showing a high degree of perturbation in the Jα-helix. Intact residues are highlighted in magenta, non-intact residues are in cyan, completely broadened residues are in blue, and unassigned residues are in black. FMN is shown in green stick.

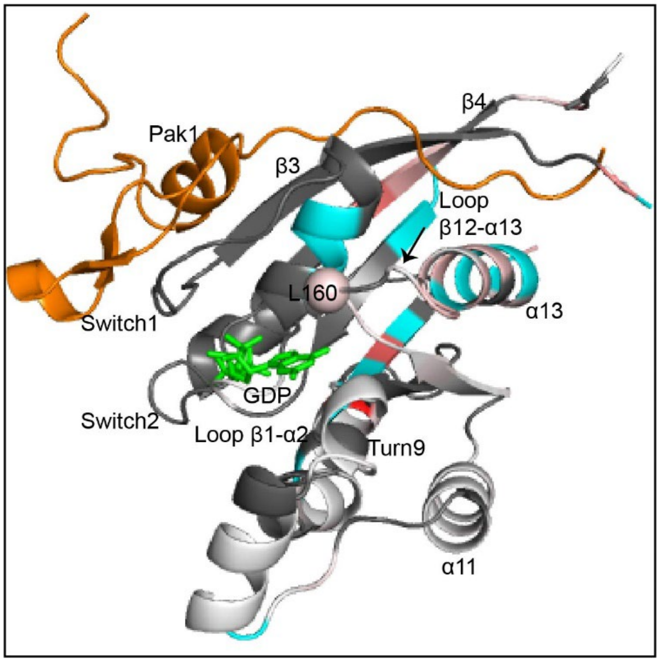


**Figure S6.** Perturbations to the nucleotide and effector binding sites. CSPs are shown in white-to-red scale. Residues highly perturbed are in red, whereas less perturbed residues are shown in white. In cyan are the broadened residue. Unassigned residues are black. Bound GDP is shown in green sticks, and Pak1 (pdb: 1e0a) is in orange. Perturbed L160 upon I539E lit mimic mutation is shown as a sphere.

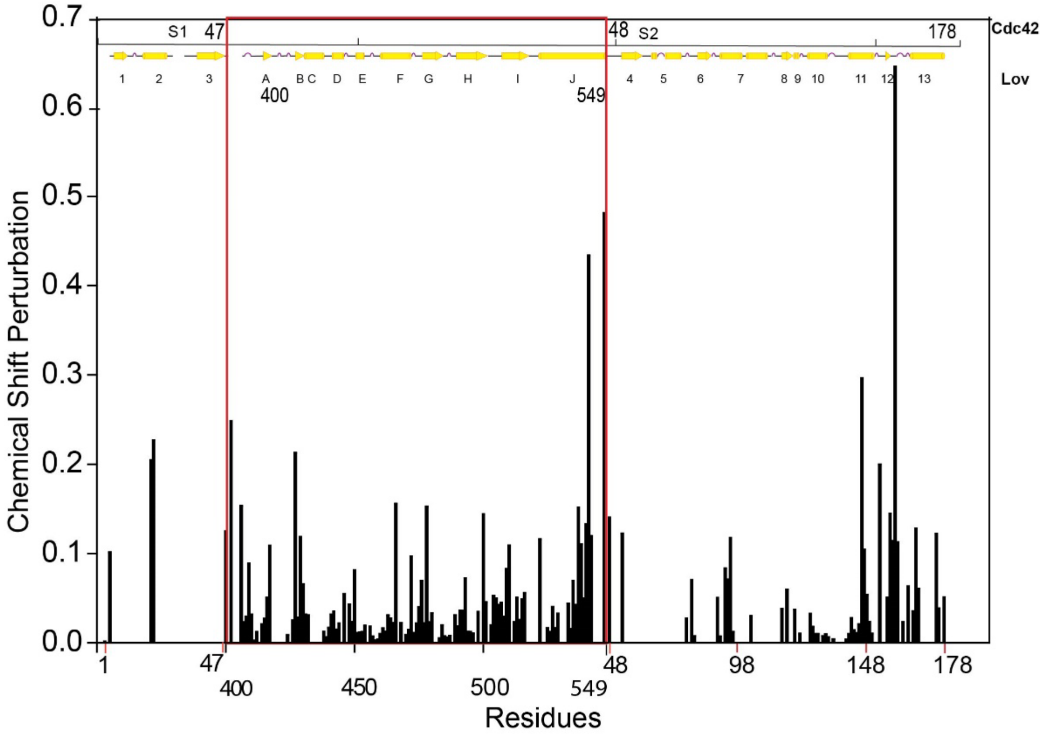


**Figure S7.** CSPs for Pak1 binding to Cdc42Lov$^{WT,CA}$.

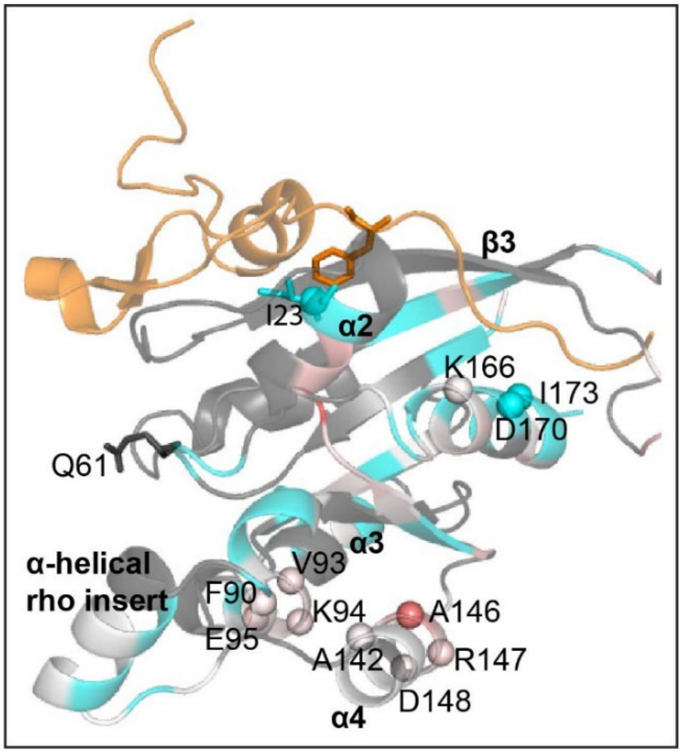


**Figure S8.** CSPs for Pak1 binding in the Cdc42 domain. Perturbations are shown in white-to-red scale. Residues highly perturbed are in red, whereas less perturbed residues are shown in white. In cyan are the broadened residue. Unassigned residues are black. Pak1 (pdb: 1e0a) is shown in orange. Key residues that show perturbation around the Pak1 binding interface are shown as spheres.

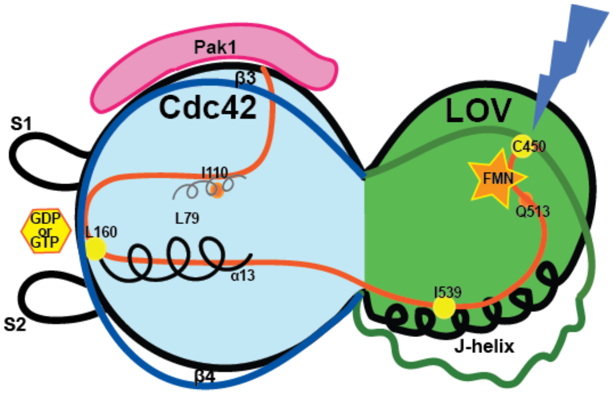


**Figure S9.** Possible allosteric pathway

#### Supplementary References

1. O. Dagliyan *et al.*, Engineering extrinsic disorder to control protein activity in living cells. *Science* **354**, 1441-1444 (2016).
